# Macrophages facilitate interclonal cooperation-induced tumor heterogeneity and malignancy by activating the innate immune signaling

**DOI:** 10.1101/2024.06.27.600955

**Authors:** Sihua Zhao, Yifan Guo, Xiaoyu Kuang, Xiaoqin Li, Chenxi Wu, Peng Lin, Qi Xie, Du Kong, Xianjue Ma

## Abstract

Tumor heterogeneity is a common hallmark of cancer and is considered a major cause of treatment failure and relapse, yet it remains poorly understood how various types of cells communicate within the tumor microenvironment (TME) to regulate tumor progression *in vivo*. Here we establish a tumor heterogeneity model in *Drosophila* eye epithelium by mutating the tricellular junction protein *M6* in cells surrounding *Ras^V12^*benign tumors and dissect the *in vivo* mechanisms underlying interclonal cooperation-induced malignancy by utilizing sophisticated genetic techniques in conjunction with single-cell RNA sequencing (scRNA-seq). Our findings reveal that loss of *M6* facilitates the malignant transformation of neighboring *Ras^V12^* tumors by activating the Toll signaling, the innate immune response pathway. Notably, inhibiting Toll signaling impedes tumor progression, whereas its activation synergistically promotes *Ras^V12^* tumor malignancy by inactivating the Hippo pathway. Mechanistically, *Ras^V12^* tumors surrounded by *M6* mutant clones lead to increased recruitment of hemocytes, which are the equivalent of macrophages in *Drosophila*, in a JNK pathway-dependent manner. Consequently, these tumor-associated macrophages secrete the Spatzle (Spz) ligand, which subsequently activates the Toll receptor within the *Ras^V12^* tumors, thereby triggering tumorigenesis. In summary, our study elucidates the complex *in vivo* interactions between genetically distinct oncogenic cells and between tumors and macrophages, shedding light on how macrophages exploit the innate immune signaling within tumors to regulate tumor heterogeneity and promote tumor progression.

**Significance statement:** Intratumoral heterogeneity profoundly affects cancer development and treatment in human tumors. The intricate nature of tumor cells and the presence of diverse cell types pose challenges to uncovering *in vivo* mechanisms responsible for heterogeneity. Our *Drosophila* tumor heterogeneity model reveals that fruit fly macrophages promotes both tumor heterogeneity and malignancy. Following recruitment by tumor cells, these macrophages secrete the ligand Spz to activate the Toll signaling pathway within tumor cells, which subsequently inactivates the Hippo pathway to drive tumorigenesis. Our study highlights the crucial role of hemocytes as intermediaries in coordinating tumor heterogeneity and facilitating intercellular communication between different cells within the TME.

## Introduction

Cancer is a multifaceted disease, with tumor heterogeneity being a critical factor that influences treatment outcomes and resistance development. This heterogeneity, characterized by the coexistence of genetically distinct subclones within a single tumor, has been increasingly recognized through the use of single-cell sequencing (sc-seq) techniques (*1–4*). These techniques have unveiled the presence of diverse cell populations within tumors and underscored the essential role of intercellular communications for tumor progression. Convincing evidence supporting this concept has also been obtained from studies utilizing mouse xenograft models, indicating that the cooperation between different subclones can lead to tumor formation (*5–8*). However, current technologies may not be sufficiently advanced to fully capture and understand the *in vivo* molecular mechanisms of tumor heterogeneity. For instance, the widely used xenograft model fails to faithfully recapitulate the TME due to its foreign and immune-compromised nature (*9*). Additionally, while sc-seq has played a crucial role in unveiling the variations among individual tumor clones, conducting *in vivo* validations subsequently poses a challenge (*10*). To overcome these limitations and comprehensively unravel the mechanisms of intercellular communication during cancer progression, an effective strategy to consider is the integration of tumor heterogeneity models that are genetically traceable and editable with the sc-seq technique.

The validity of *Drosophila* as a model organism in cancer biology has been well established, with numerous cancer-related signaling pathways first identified in this organism (e.g., Notch, Wnt, Hedgehog and Hippo pathway) (*11, 12*). The powerful genetic tools available in *Drosophila* make it an optimal model system for elucidating the mechanisms of oncogenic intercellular communications (*13–17*). Accumulating evidence suggests that *Drosophila* genetic models faithfully replicate *in vivo* oncogenic cell-cell interactions, as clones harboring distinct oncogenic mutations can collaboratively promote tumor progression (*18, 19*). For example, oncogenic *Ras* (*Ras^V12^*) clones with defects in the mitochondrial respiratory complex could lead to the malignancy of neighboring benign *Ras^V12^* tumors through the induction of Unpaired (Upd, an IL-6 homolog) and Wingless (Wg, a Wnt homolog) (*20*). Likewise, benign *Ras^V12^* tumors exhibit a shift towards malignancy upon being encompassed by clones featuring overexpression of the oncoprotein *Src* or clones that are deficient in polarity due to *scribble* (*scrib*) mutations (*21, 22*).

Tricellular junction (TCJ) proteins are specialized proteins that are located at the points where three epithelial or endothelial cells meet. They are crucial for maintaining the integrity and function of tissues, and dysregulation of these proteins can lead to various pathological conditions, including deafness and certain types of cancer (*23, 24*). In *Drosophila*, the key TCJ proteins include Anakonda/bark beetle (Aka/bark), Gliotactin (Gli), sidekick (sdk), and M6. These proteins regulate a range of biological activities, such as disengagement of daughter cells during cytokinesis, TCJ assembly, positioning of photoreceptor neurons, and cell contraction (*25–28*). Interestingly, mutations in *M6* could synergize with *Ras^V12^*to drive cell-autonomous overgrowth and apical cell delamination (*29*). However, it remains unknown whether *M6* contributes to tumor heterogeneity.

In this study, we combine the genetic tractability of *Drosophila* with the power of single-cell sequencing for an in-depth dissection of tumor heterogeneity. Using the *Drosophila* eye imaginal epithelium, we discovered that clones bearing *M6* mutations could promote the tumor progression of neighboring benign *Ras^V12^*tumors. Our bulk RNA-seq and single-cell RNA sequencing (scRNA-seq) analyses revealed that subclonal cooperation-induced malignancy was due to the activation of the innate immune-responding Toll pathway and subsequent inactivation of the Hippo pathway. Notably, we revealed the specific upregulation of the Toll ligand, Spz, in tumor associated macrophages. Moreover, we demonstrated that macrophage derived Spz is both necessary and sufficient for the malignant transformation of *Ras^V12^* tumors. In summary, our findings shed light on a previously unrecognized role of the Toll pathway and macrophages in promoting tumorigenesis, while providing a mechanistic understanding of how macrophages regulate epithelial tumor heterogeneity by hijacking the innate immune system.

## Results

### Interclonal cooperation between *Ras^V12^* and *M6^-/-^* promotes tumor malignancy

To establish a tumor heterogeneity model in the eye-antennal imaginal discs of *Drosophila* larvae, we modified the widely used genetic recombinase system known as MARCM (mosaic analysis with a repressible cell marker), which was originally designed to generate positively labeled homozygous mutant clones (*30*). We adapted this system by recombing the mutations of interest (*X*) with the corresponding Gal80 onto the same chromosome arm (Fig. S1A). This design allows for the generation of two neighboring daughter cells with different genotypes after mitosis: one GFP negative clone carrying the homozygous mutant of *X*, and one GFP positive clone with ectopic *Ras^V12^* expression (Figs. 1A and S1A). *M6* mutant clones undergo cell-autonomous apoptosis and are eliminated, at least partially, through a caspase-dependent mechanism (Figs. S1B and B’), whereas clonal expression of *Ras^V12^* alone induces hyperproliferation and forms benign tumors (Figs. 1B-E). Remarkably, the deletion of *M6* in *Ras^V12^*surrounding cells transforms GFP^+^ *Ras^V12^* clones into malignant tumors (Figs. 1B-E), indicating that interclonal cooperation between *Ras^V12^*and *M6* (referred to as *Ras^V12^//M6^-/-^*) drives tumor malignancy. Moreover, despite the hyperproliferative nature of *Ras^V12^* cells, regional caspase activation was observed in *Ras*^V12^ tumors surround by *M6* clones (Figs. S1C and C’), in accordance with the previously reported tumor-promoting role of caspase activation (*31*).

**Figure 1.**
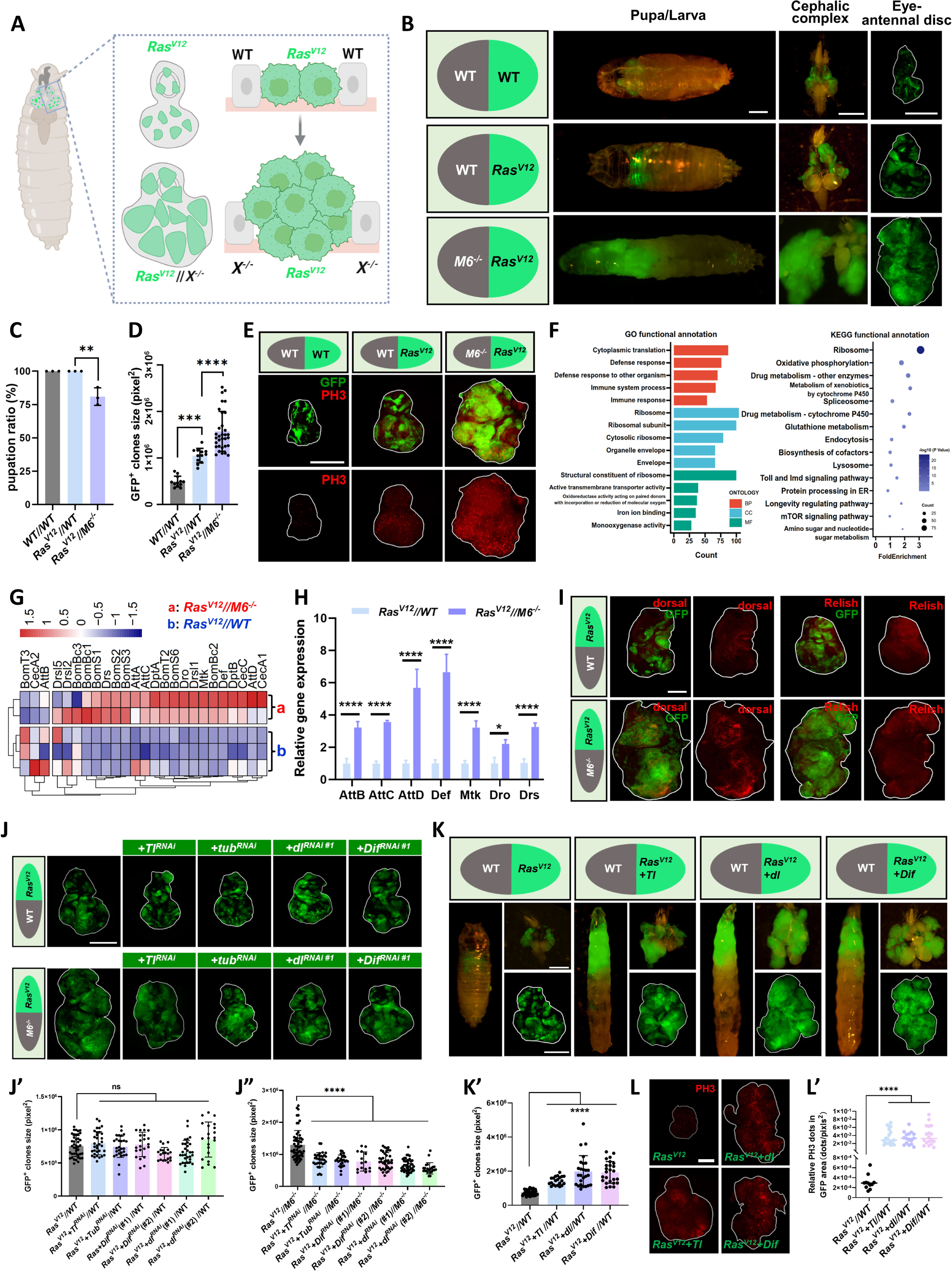
Interclonal cooperation between *Ras^V12^* and *M6^-/-^* promotes tumor malignancy by activating the Toll pathway. **(A)** A cartoon illustrating the strategy of generating heterogeneity tumor model in *Drosophila*. **(B)** Late-stage larva/pupa (left), cephalic complex (middle) and eye-antennal disc (right) bearing GFP-labeled wild-type clones (green)//non-GFP-labeled wide-type clones (*WT//WT*), *Ras^V12^*-expressing clones (green)//wide-type clones (*Ras^V12^//WT*), and *Ras^V12^*-expressing clones (green)//*M6* mutant clones (*Ras^V12^//M6^-/-^*). **(C)** Quantification of pupation ratio of larvae with indicated genotypes: *WT//WT*, *Ras^V12^//WT*, *Ras^V12^//M6*^-/-^. **(D)** Quantification of GFP^+^ clones’ size with indicated genotypes (from left to right, n = 12, 12, 32). **(E)** Eye-antennal discs bearing clones of *WT//WT*, *Ras^V12^//WT* and *Ras^V12^//M6^-/-^* were stained with anti-PH3 antibody. **(F)** The top five enriched terms of the GO regulon and top 15 enriched KEGG terms obtained from 1504 up-regulated genes by comparing the *Ras^V12^//M6^-/-^* with *Ras^V12^//WT* group. **(G)** Heatmap of normalized Toll and Imd signaling target gene expression between the *Ras^V12^//M6^-/-^* and *Ras^V12^//WT* samples. Normalized gene expression of heatmap was calculated as log2(CPM+1) and scaled by row. **(H)** qPCR to determine relative AMPs expression level of *Ras^V12^*//WT and *Ras^V12^//M6^-/-^* tumors. **(I)** Eye-antennal discs bearing clones of WT//WT, *Ras^V12^*//WT, and *Ras^V12^//M6^-/-^* were stained with dorsal (left) and Relish (right) antibodies. **(J)** Eye-antennal discs bearing *Ras^V12^//WT*, *Ras^V12^+ Tl^RNAi^//WT*, *Ras^V12^+ Tub^RNAi^//WT*, *Ras^V12^+ dl^RNAi#1^//WT*, *Ras^V12^+ Dif^RNAi#1^//WT*, *Ras^V12^//M6^-/-^*, *Ras^V12^+ Tl^RNAi^//M6^-/-^*, *Ras^V12^+ Tub^RNAi^//M6^-/-^*, *Ras^V12^+ dl^RNAi#1^//M6^-/-^*, and *Ras^V12^+ Dif^RNAi#1^//M6^-/-^*. Quantification of GFP^+^ clones’ size for the indicated genotypes (J’, from left to right, n = 42, 33, 33, 28, 20, 31, 21) and (J”, from left to right, n = 60, 30, 26, 15, 39, 44, 23). **(K)** Late-stage larva/pupa, cephalic complex and eye-antennal discs bearing *Ras^V12^//WT*, *Ras^V12^+ Tl//WT*, *Ras^V12^+ dl//WT*, *Ras^V12^+ Dif//WT*. Quantification of GFP^+^ clones’ size (K’, from left to right, n = 42, 21, 26, 25). **(L)** Eye-antennal discs bearing clones of *Ras^V12^//WT*, *Ras^V12^+ Tl//WT*, *Ras^V12^+ dl//WT*, *Ras^V12^+ Dif//WT* were stained with anti-PH3 antibody. Quantification of PH3 dots per GFP^+^ clones’ area (L’, from left to right, n=11, 19, 18, 18). Statistical analysis by Šídák’s multiple comparisons test (H) and Tukey’s multiple comparisons tests (C, D, J’, J’’, K’, L’); mean ± SD. ns., not significant; \**p* < 0.05, ** *p* < 0.01, \*\*\**p* < 0.001, \*\*\*\**p* < 0.0001. Scale bars: 400μm (B, E, K), 200μm (I, J, L).

### *Ras* synergizes with Toll signaling to promote malignancy in *Ras^V12^//M6^-/-^* tumors

Next, to dissect the mechanism that regulates the malignant transformation of *Ras^V12^* tumors, we performed bulk RNA-sequencing analysis on both *Ras^V12^* tumors and *Ras^V12^//M6^-/-^*tumors. The Gene Ontology (GO) and Kyoto Encyclopedia of Genes and Genomes (KEGG) analyses against 1504 up-regulated genes revealed a significant enrichment of defense response, immune response and immune related signaling pathways (Fig. 1F). In *Drosophila*, there are two major innate immunoregulatory pathways known as the Toll pathway and the immune deficiency (Imd) pathway, they collectively control the systemic production of antimicrobial peptides (AMPs) to combat microbial infection (*32–34*). We observed a marked increase in multiple AMPs in the *Ras^V12^//M6^-/-^*tumors (Fig. 1G), which was subsequently confirmed through qRT-PCR analysis (Fig. 1H). Interestingly, a strong upregulation of dorsal (dl), the downstream transcription factor of the Toll pathway, was observed, while there was no change in the expression of Relish (Rel), the downstream transcription factor of the Imd pathway (Fig. 1I), indicating that Toll pathway might be more important in *Ras^V12^//M6^-/-^*tumors. The Toll pathway can be canonically activated by Gram-positive bacterial infections and fungal infections (*32, 35*). To explore whether the activation of Toll signaling induced by *Ras^V12^//M6^-/-^*is a consequence of infection, we utilized axenic cultures and antibiotic cocktail treatment to deplete the bacterial microbiome to below detectable levels (Fig. S1D). Interestingly, the elimination of bacteria further enhanced the overgrowth of *Ras^V12^//M6^-/-^*tumor (Figs. S1E-F), suggesting a potential growth-inhibition role of bacteria in *Drosophila*. Additionally, we noticed that despite a significant reduction in AMP expression after antibiotic treatment, *Ras^V12^//M6^-/-^* tumor still exhibited higher AMP expression levels compared to *Ras^V12^* tumor (Figs. S1G), implying that the interclonal cooperation between *Ras^V12^* and *M6^-/-^* can induce Toll activation independently of infection.

We next investigated whether the Toll pathway activation is essential for the malignant transformation of *Ras^V12^* tumors. Inhibition of multiple components of Toll signaling, such as *Toll* (*Tl*), *tube* (*tub*), *dl*, or *Dorsal-related immunity factor* (*Dif*), did not notably affect *Ras^V12^*-induced benign tumorous growth (Figs. 1J and 1J’). However, these interventions significantly impeded *Ras^V12^//M6^-/-^*-induced tumor progression (Figs. 1J and 1J”). Remarkably, genetically activating the Toll pathway by co-expressing an activated form of *Tl* or the transcription factors *dl* or *Dif* could phenocopy that of *Ras^V12^//M6^-/-^* tumor and transformed *Ras^V12^* into malignant tumors (Figs. 1K-L’ and S1H-H”). These findings collectively suggest that the activation of the Toll pathway is both necessary and sufficient for the malignant transformation of *Ras^V12^*tumors.

### *Ras^V12^*//*M6^-/-^* promotes tumor malignancy through Toll pathway-mediated inactivation of Hippo signaling

To further elucidate the molecular mechanisms underlying tumor malignancy driven by interclonal communications in *Ras^V12^//M6^-/-^*tumor, we performed scRNA seq analysis on eye-antennal discs dissected from wild type (WT) and *Ras^V12^//M6^-/-^*larvae (Fig. 2A). Based on the similarities in the gene expression profiles of high-quality cells, 32225 cells from WT and 43992 cells from *Ras^V12^//M6^-/-^*tumors were collected and assembled into 27 clusters that were categorized into three groups: epithelium cells, hemocytes, and glial cells (Figs. S2A and S2B). Unique clusters to the *Ras^V12^//M6^-/-^* tumors were separated and re-clustered (Figs. S2C and S2D), and epithelium cells were then isolated for further analyses (Fig. 2B). According to the expression profiles of exogenously expressed GFP, these 19 clusters were categorized into two types. Clusters that demonstrated a high level of occupancy, with over 65% of cells expressing GFP, were identified as GFP-positive clusters, while the remaining ones were classified as GFP-negative clusters (Fig. 2C). In line with our data on Toll pathway activation, several GFP-positive clusters display elevated *dl* expression (Fig. 2D). Cells derived from GFP-positive clusters were subsequently isolated and integrated with WT epithelium cells, further reassembled into 19 individual clusters (Fig. 2E), and 5 clusters that are unique to the *Ras^V12^//M6^-/-^*tumors were separated (Fig. 2F). Subsequently, we performed KEGG pathway analysis on differentially expressed genes (DEGs) within each cluster and identified several overlapping regulons across different clusters (Figs. 2G and S2E). Apart from the Toll and Imd signaling, we noticed an enrichment of Hippo signaling (Fig. 2G), a master regulator of tissue growth and tumorigenesis (*36–41*). Furthermore, upon re-analysis of the DEGs derived from bulk RNA-seq data of *Ras^V12^*and *Ras^V12^//M6^-/-^* tumors, we observed that Hippo signaling is also significantly enriched (Fig. 2H).

**Figure 2.**
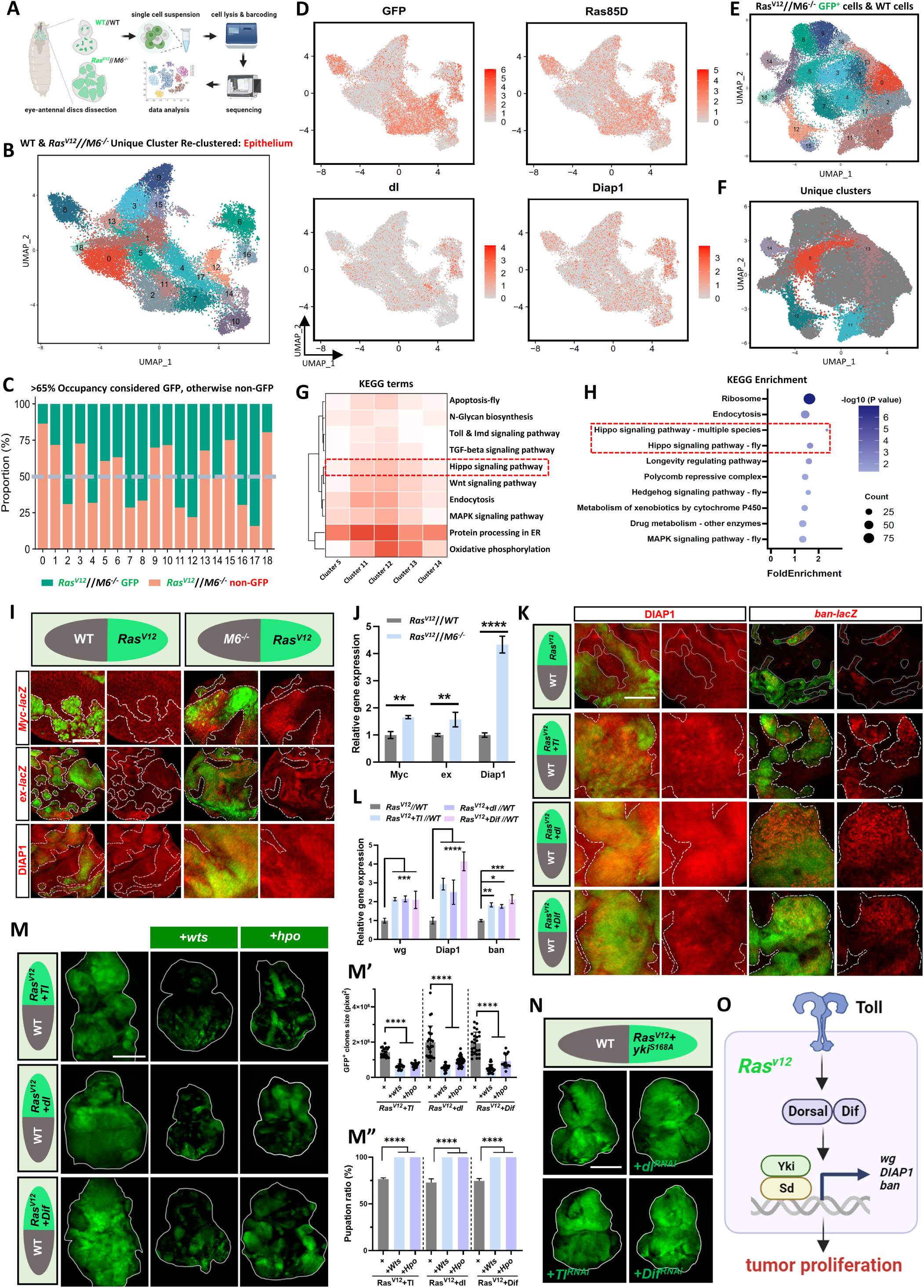
*Ras^V12^*//*M6^-/-^* promotes tumor malignancy through Toll pathway-mediated inactivation of Hippo signaling. **(A)** Scheme of the experimental workflow for preparing *Drosophila* tumor samples and conducting subsequent single-cell RNA-seq analysis. **(B)** Uniform manifold approximation and projection (UMAP) plot showing major cell types of re-clustered *Ras^V12^//M6^-/-^* unique epithelial cells. Dots represent individual cells, and colors represent distinct cell populations. A total of 19 clusters were generated (resolution = 1.3). **(C)** Bar plot showing the fraction of GFP-positive cells and non-GFP cells in each cluster. Clusters having at least 330 GFP^+^ cells and over 65% GFP^+^ cell proportion were considered as GFP^+^-dominant clusters. **(D)** UMAP plots showing the marker genes expression of *GFP*, *Ras85D*, *dl* (*dorsal*), and *Diap1* for the *Ras^V12^//M6^-/-^* unique epithelial cells. **(E)** UMAP plot showing the integration of *Ras^V12^//M6^-/-^*GFP-positive epithelial cells and WT epithelial cells. A total of 19 clusters were generated (resolution = 1.2). **(F)** UMAP plot showing clusters unique to the *Ras^V12^//M6^-/-^*GFP^+^ cells compared to WT cells, including cluster 5, 11, 12, 13, and 14. **(G)** Eight overlapped enriched KEGG terms between clusters 5, 11, 12, 13, and 14 are shown, together with additional terms that were specifically enriched in specific clusters. **(H)** KEGG enrichment of all DEGs from bulk RNA-seq between *Ras^V12^//M6^-/-^* and *Ras^V12^//WT*. **(I)** *Myc-lacZ*, *ex-lacZ*, and DIAP1 staining in eye discs bearing *Ras^V12^//WT* and *Ras^V12^//M6^-/-^*. **(J)** qPCR analysis to determine the mRNA levels of *yki* target genes (*Myc*, *ex*, and *Diap1*) in eye-antennal discs of indicated flies. **(K)** Eye-antennal discs bearing clones of *Ras^V12^//WT*, *Ras^V12^+ Tl//WT*, *Ras^V12^+ dl//WT*, *Ras^V12^+ Dif//WT* were stained with anti-DIAP1 and anti-β-galactosidase antibodies for the *ban-lacZ*. **(L)** qPCR analysis to determine the mRNA levels of *yki* target genes (*wg*, *Diap1*, and *ban*) in eye-antennal discs of indicated flies. **(M)** Eye-antennal discs bearing clones of *Ras^V12^*+*Tl//WT*, *Ras^V12^*+*Tl*+*wts//WT*, *Ras^V12^*+*Tl*+ *hpo//WT*, *Ras^V12^*+*dl//WT*, *Ras^V12^*+*dl*+*wts//WT*, *Ras^V12^*+*dl*+*hpo//WT*, *Ras^V12^+Dif//WT*, *Ras^V12^*+*Dif*+*wts//WT*, *Ras^V12^*+*Dif*+*hpo//WT*. Quantification of GFP^+^ clones’ size (M’, from left to right, n = 21, 22, 20, 26, 22, 38, 25, 22, 16) and pupation ratio (M”) of flies with indicated genotypes. **(N)** Eye-antennal discs bearing clones of *Ras^V12^*+*yki^S168A^//WT*, *Ras^V12^*+*yki^S168A^*+*Tl^RNAi^//WT*, *Ras^V12^*+*yki^S168A^*+*dl^RNAi^//WT*, *Ras^V12^*+*yki^S168A^*+*Dif^RNAi^//WT*. **(O)** A model shows that Toll pathway activation in *Ras^V12^* clones promotes malignancy through transcriptional upregulation of Yki/Sd target genes. Statistical analysis by Šídák’s multiple comparisons test (J and L) or Tukey’s multiple comparisons test (M’ and M”); mean ± SD. \**p* < 0.05, \*\**p* < 0.01, \*\*\**p* < 0.001, \*\*\*\**p* < 0.0001. Scale bars: 100μm (I, K), 400μm (M, N).

Next, we investigated the role of Hippo signaling in *Ras^V12^//M6^-/-^*-induced tumor progression. *Ras^V12^//M6^-/-^*tumors displayed significant upregulation of multiple Hippo pathway target genes, including *Myc*, *expanded* (*ex*), and *Death-associated inhibitor of apoptosis 1* (*Diap1*) (Fig. 2I), as confirmed by qRT-PCR analysis (Fig. 2J) and scRNA-seq analysis (Fig. 2D). The key components of Hippo pathway consist of *hippo* (*hpo*), *warts* (*wts*), and *yorkie* (*yki*) (reviewed in (*^42^*)). Hpo phosphorylates Wts, which subsequently phosphorylates and inactivates Yki. We observed that inhibiting Yki activity through either the removal of one copy of endogenous *yki* or ectopic expression of *wts* significantly suppressed the overgrowth of *Ras^V12^//M6^-/-^* tumors (Figs. S2F and S2F’). We further dissected the genetic interactions between Toll and Hippo pathways *in vivo*. Our findings demonstrated that *Ras^V12^*tumors with increased Toll pathway activity markedly increased the expression of Hippo pathway target genes (Figs. 2K, 2L, and S2G). Consistently, inhibiting Yki activity by co-expressing *wts* or *hpo* significantly impeded the synergistic tumor-promoting effect between *Ras^V12^* and Toll activation (Fig. 2M and 2M’), and restored the pupation defects (Fig. 2M”). Conversely, inhibiting Toll activity did not influence the overgrowth phenotype induced by co-expressing *Ras^V12^* and *Yki^S168A^* (Figs. 2N and S2H). Together, these genetic data demonstrate that Toll pathway activation promotes malignancy by inhibiting the Hippo pathway in *Ras^V12^//M6^-/-^*tumors (Fig. 2O).

### Macrophage derived Spz activates Toll pathway to facilitate malignant transformation of *Ras^V12^* tumors

Next, we investigated the mechanism by which Toll pathway is activated. Spz is a secreted cytokine and acts as the major ligand for the Toll receptor in *Drosophila*, it plays a crucial role in the immune response and development (*32, 35*). Surprisingly, in contrast to *Ras^V12^* benign tumors, *Ras^V12^//M6^-/-^* tumors exhibited a remarkable increase in the expression of Spz in the non-GFP regions (Fig. 3A). Upon closer examination, we noticed that Spz specifically accumulated in cells surrounding *Ras^V12^* tumors (Fig. 3A). In contrast, no significant changes in *spz* expression were detected in *Ras^V12^//M6^-/-^* tumors or *M6* mutated clones (Figs. S3A and S3B). To examine the role of *spz* in the tumor progression of *Ras^V12^//M6^-/-^*, *spz* was removed from the entire larvae. Remarkably, this resulted in a significant reduction in tumor size (Fig. 3B), accompanied by the inhibition of upregulation of dl and target genes of the Hippo pathways (Figs. 3C and S3C). Importantly, the re-activation of the Toll pathway in the genetic background lacking *spz* effectively reversed the tumor suppression phenotype (Fig. 3B), suggesting that Spz plays a crucial role in the activation of Toll pathway within the *Ras^V12^* tumors, thereby facilitating their malignant transformation.

**Figure 3.**
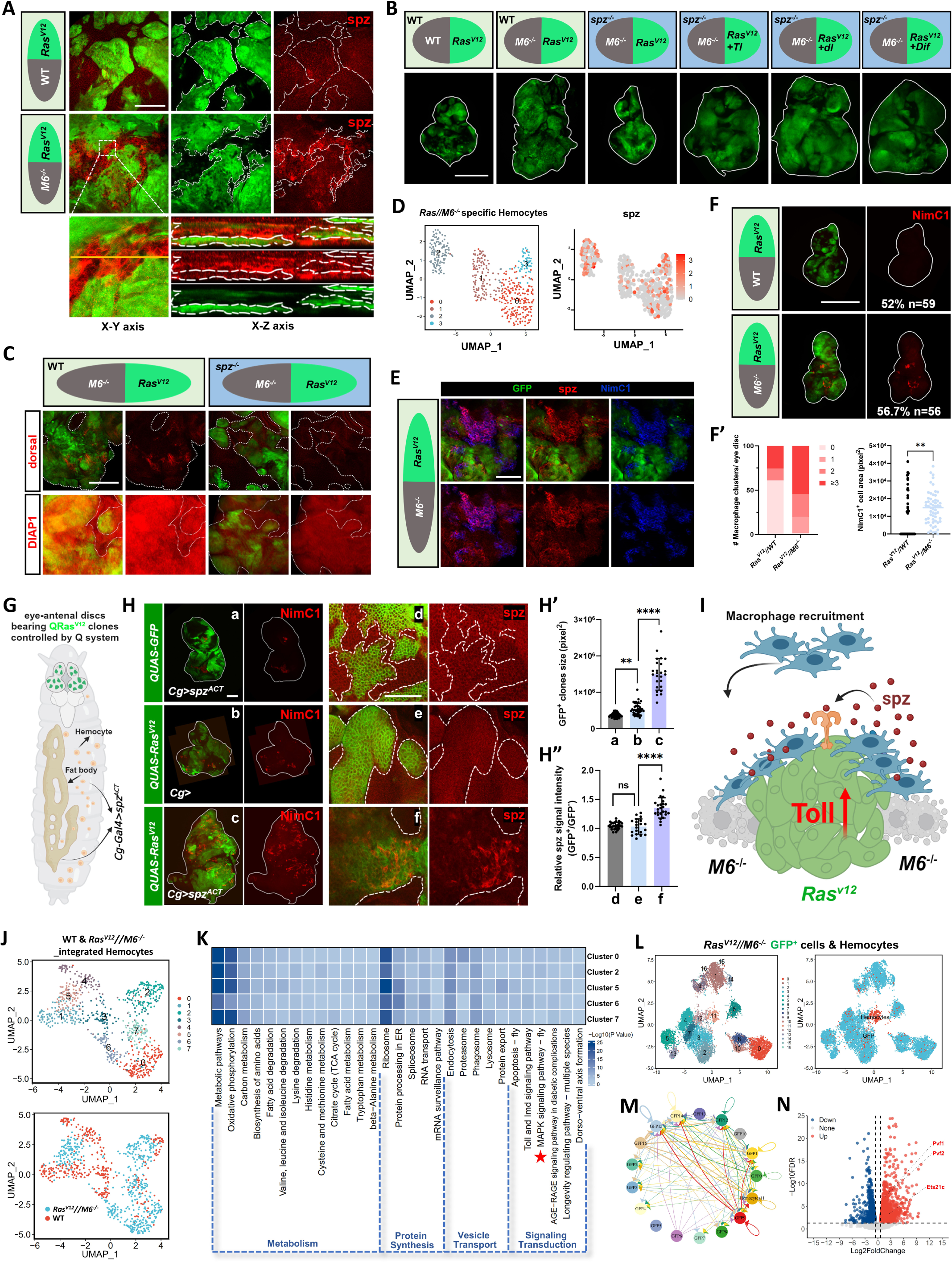
Macrophage derived Spz activates Toll pathway to facilitate malignant transformation of *Ras^V12^* tumors. **(A)** Eye discs bearing clones of *Ras^V12^//WT* and *Ras^V12^//M6^-/-^* were stained with anti-spz antibody. Bottom images show xy cross-section and xz across-section. Lines represent the position of lateral section images. **(B)** Eye-antennal discs bearing clones of *Ras^V12^//WT*, *Ras^V12^//M6^-/-^*, *Ras^V12^*+*spz^-/-^//M6^-/-^*+*spz^-/-^*, *Ras^V12^*+*Tl*+*spz^-/-^*//*M6^-/-^*+*spz^-/-^*, *Ras^V12^*+*dl*+*spz^-/-^//M6^-/-^*+*spz^-/-^*, *Ras^V12^*+*Dif*+*spz^-/-^//M6^-/-^*+*spz^-/-^*. **(C)** Eye discs bearing clones of *Ras^V12^//M6^-/-^* and *Ras^V12^*+*spz^-/-^//M6^-/-^+spz^-/-^*were stained with anti-dorsal and anti-DIAP1 antibody. **(D)** UMAP plots showing re-clustered *Ras^V12^//M6^-/-^* unique macrophages (hemocytes) and *spz* expression. **(E)** Eye disc bearing clones of *Ras^V12^//M6^-/-^* were stained with both anti-spz and anti-NimC1 antibodies. **(F)** Eye-antennal discs bearing clones of *Ras^V12^//WT* and *Ras^V12^//M6^-/-^* were stained with anti-NimC1 antibody, and the phenotype penetrance are shown. (F’) Quantification of the number of macrophages clusters and area (from left to right, n = 59, 56). **(G)** Cartoon illustrating the dual expression system to overexpress activated *spz* (*spz^ACT^*) in the hemocytes and fat bodies under the *Cg* promoter and simultaneously inducing GFP labeled *QRas^V12^* clones in the eye-antennal disc controlled by the Q system. **(H)** Eye-antennal discs bearing GFP labeled clones of wild type with *spz^ACT^*overexpression by the *Cg* promoter (a, d) and *QRas^V12^* overexpression clones without (b, e) or with (c, f) *spz^ACT^* expression by the *Cg* promoter were stained with anti-NimC1 or anti-spz antibody. Quantification of GFP^+^ clones’ size (H’, from left to right, n = 48, 39, 24) and relative spz signal intensity (H”, from left to right, n = 28, 20, 25) of indicated flies. **(I)** A model shows that tumors-associated macrophages secrete spz to activate the Toll pathway, leading to the malignancy of *Ras^V12^* tumors. **(J)** UMAP plots of integrated WT and *Ras^V12^//M6^-/-^* tumor-derived hemocytes showing origin-specific clusters. **(K)** Heatmap showing the enriched 29 overlapping KEGG terms in the 5 clusters (0, 2, 5, 6, 7) of *Ras^V12^//M6^-/-^* tumor-derived hemocytes. **(L)** UMAP plots of integrated *Ras^V12^//M6^-/-^*tumor-derived hemocytes and *Ras^V12^//M6^-/-^* GFP^+^ tumor cells. A total of 17 clusters were identified (left, resolution = 1.2). **(M)** Circle plot showing a significant enrichment in the PVR signaling pathway between multiple *Ras^V12^//M6^-/-^*GFP^+^ tumor cell clusters and *Ras^V12^//M6^-/-^*tumor-derived hemocytes (marked with an asterisk). **(N)** Volcano plot of bulk RNA-seq data by comparing *Ras^V12^//M6^-/-^*malignant tumors with *Ras^V12^*//WT benign tumors. Statistical analysis by Tukey’s multiple comparisons test (H’, H”) or Mann Whitney test (F’); mean ± SD. ns., not significant; \*\**p* < 0.01, **** *p* < 0.0001. Scale bars: 50μm (A, E), 100μm (C, H), and 400μm (B, F).

One of the primary sites for the production and activation of the Spz protein in *Drosophila* is the hemocytes (blood cells)(*43, 44*). Among these cells, over 90-95% are composed of plasmatocytes, which are the equivalent of mammalian macrophages (*45*). In line with this, our analysis of scRNA-seq data also revealed the presence of multiple clusters of hemocytes (Fig. S2A), leading us to hypothesize that macrophages could serve as the source of Spz. Indeed, scRNA-seq analysis revealed high expression of *spz* in hemocytes (Fig. 3D). Moreover, by utilizing antibody staining against NimC1, a widely accepted marker for *Drosophila* macrophages (*46*), we found that tumor-associated macrophages (TAMs) exhibit increased expression of Spz (Fig. 3E). We also observed a significant increase in the number of TAM in *Ras^V12^//M6^-/-^*tumors (Figs. 3F and 3F’). To further confirm that hemocytes are the source of Spz, we combined the Q MARCM system with Gal4/UAS system to examine whether macrophage-derived Spz could promote *Ras^V12^* tumor transformation (Fig. 3G). Notably, the ectopic expression of an activated form of *spz* (*spz^ACT^*) under the control of *Cg-Gal4* (which labels both hemocytes and fat bodies) remotely accelerated *QRas^V12^* tumor overgrowth and macrophage recruitment (Figs. 3H-H”) and upregulated dl and the Hippo pathway target gene expression (Fig. S3D). However, overexpression of *spz^ACT^*within the *Ras^V12^* clones had no effect on tumor size (Figs. S3E and S3E’). Together, these data suggest that macrophage derived Spz activates Toll pathway and subsequently transforms *Ras^V12^* clones into malignant tumors (Fig. 3I).

### Systemic analysis of TAM and tumor-macrophage intercellular communication

To explore the potential mechanisms underlying the upregulation of *spz* expression by hemocytes, we integrated hemocytes derived from both wild type and *Ras^V12^//M6^-/-^* tumor bearing discs and identified five clusters that are unique to *Ras^V12^//M6^-/-^* tumors (Figs. 3J and S3F). The DEGs between the WT hemocytes and *Ras^V12^//M6^-/-^* tumor-derived hemocytes were extracted for further functional annotation, and a total of 29 overlapping enriched regulons were identified from KEGG database, we zoomed in and discovered two positively correlated pathways, the immune pathway and MAPK (mitogen-activated protein kinase) pathway (Figs. 3K and S3G-K). *Drosophila* encompasses three main MAPK pathways: the ERK (extracellular-signal-regulated kinase), p38, and JNK (c-Jun N-terminal kinase) pathways. Interestingly, previous reports have indicated that the activation of the JNK pathway upregulates *spz* expression in *Drosophila* (*47*). In line with this, we observed a positive correlation between the expression of *spz* and the activation of JNK specifically in *Ras^V12^//M6^-/-^* tumor-derived hemocytes (Fig. S3K). Collectively, these findings indicate that the hyperactivation of JNK signaling in the hemocytes may contribute to the upregulation of *spz*.

Next, we integrated both GFP-positive and GFP-negative clusters from *Ras^V12^//M6^-/-^* tumors with hemocytes profiles (Fig. 3L and S3O) and explored the potential tumor-hemocyte communications with FlyPhoneDB, a comprehensive web-based tool for predicting cell-cell communications in *Drosophila* (*48*). By analyzing the average ligand and receptor expression values for each cluster, we identified three signaling pathways with significant interaction scores (*p* < 0.05) within GFP-positive tumor cells and hemocytes integrated datasets: the EGFR, FGFR, and PVR pathways. Among these, only the PVR pathway exhibited exceptionally active tumor-hemocyte crosstalk (Figs. 3M and S3L). The GFP-positive *Ras^V12^//M6^-/-^* tumor subclusters displayed elevated expression of the PVR ligands (*Pvf1*, *Pvf2*, and *Pvf3*) (Figs. S3L and S3M), while the hemocytes showed high levels of the receptor *Pvr* (Fig. S3N). Consistent with these findings, our bulk RNA-seq data showed significant upregulation of PVR ligands in *Ras^V12^//M6^-/-^*tumor samples (Fig. 3N). However, we did not observe any significant connectivity between GFP-negative clusters and hemocytes. Intriguingly, Pvr has also been identified as a potent activator of JNK signaling in *Drosophila* (*49*), while Pvfs are essential for hemocyte migration and proliferation (*50*). Taken together, our scRNA-seq analysis indicates that tumor secreted Pvfs might attract and activate the Pvr on the hemocytes, which subsequently activate JNK signaling and lead to the upregulation of *spz*.

### JNK activity propagates to surrounding *Ras^V12^* cells to facilitate tumorigenesis and macrophage recruitment

Next, we further investigated the mechanisms by which *Ras^V12^//M6^-/-^*tumor upregulates *Pvfs* expression. It has been previously demonstrated that the transcription factor Ets21C can directly promote the transcriptional activation of *Pvf1*(*51*). Moreover, Ets21C is also known to be a downstream target of JNK signaling in diverse contexts, including tumorigenesis, intestinal renewal, and regeneration (*51–54*). In line with these findings, our scRNA-seq data revealed that GFP-positive *Ras^V12^//M6^-/-^*tumor cells exhibit significant enrichment of the MAPK pathway (Fig. 4A), as well as elevated expression of the positive regulators (*msn*, *bsk*, *kay*, and *Jra*) and the target genes (*puc* and *Mmp1*) of the JNK pathway (Fig. S4A). Consistently, antibody staining indicated that *Ras^V12^//M6^-/-^* tumors display robustly increased expression of Mmp1 (Fig. 4B). Furthermore, the expression of *Ets21C* was also upregulated in both our scRNA-seq and bulk RNA-seq data of *Ras^V12^//M6^-/-^* tumors (Figs. 3N and S3L). We next investigated whether JNK pathway activation is required for tumor progression by expressing a dominant negative form of *bsk* (the *Drosophila* JNK, *bsk^DN^*). Inhibiting the JNK pathway dramatically impeded *Ras^V12^//M6^-/-^*-induced tumor overgrowth and Mmp1 upregulation (Figs. 4B and 4B’). More importantly, blocking JNK activation also completely abolished the recruitment of TAM (Figs. 4B and 4B”). These results collectively demonstrate that JNK pathway activation within the *Ras^V12^*tumor facilitates the recruitment of macrophages and subsequently promotes tumorigenesis.

**Figure 4.**
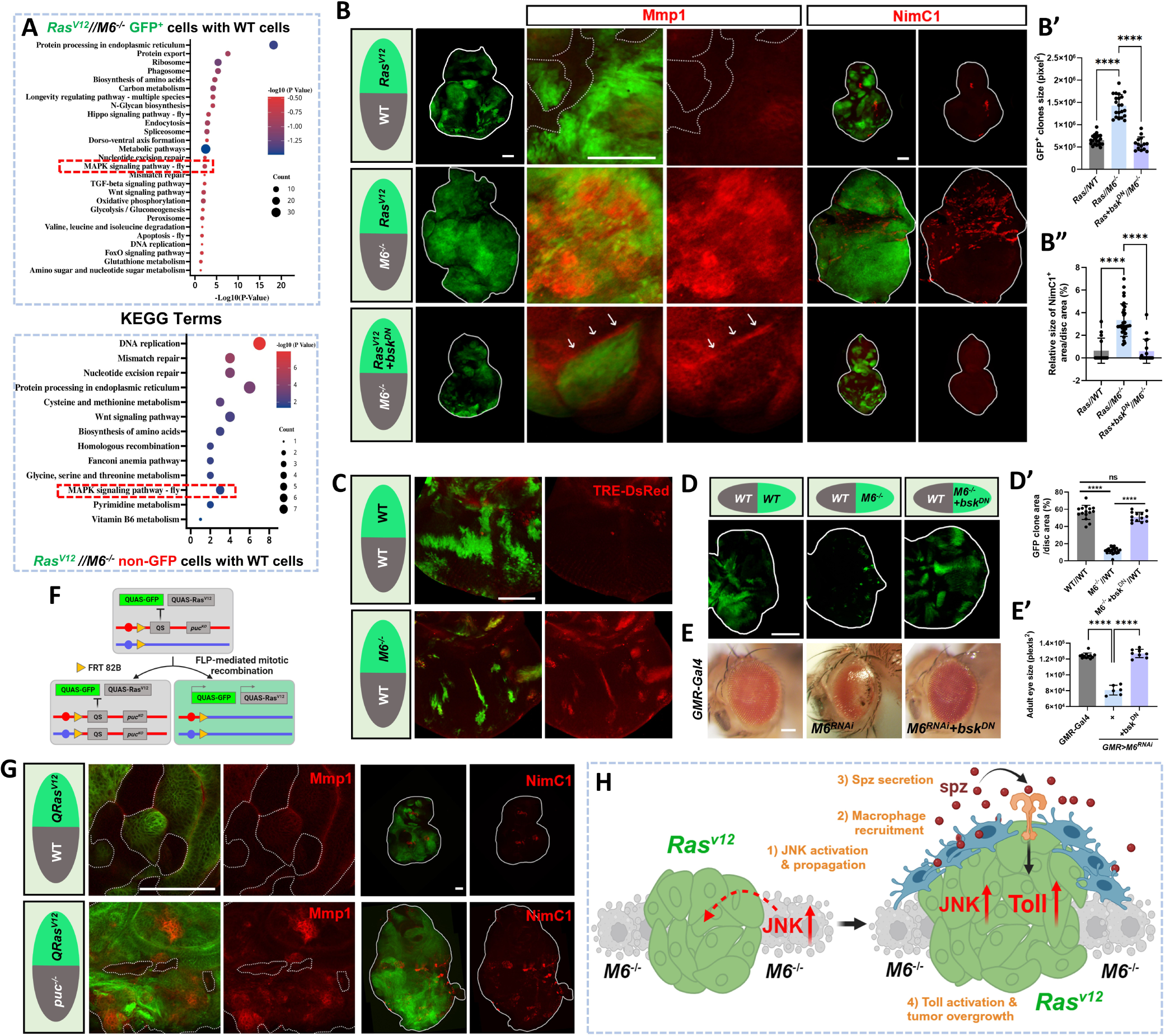
JNK activity propagates to surrounding *Ras^V12^* cells to facilitate tumorigenesis and macrophage recruitment. **(A)** KEGG enrichment analysis comparing *Ras^V12^//M6^-/-^* GFP^+^ cells to WT cells (top), and *Ras^V12^//M6^-/-^* non-GFP cells to WT cells (bottom), respectively. **(B)** Eye discs bearing clones of *Ras^V12^ //WT*, *Ras^V12^//M6^-/-^* and *Ras^V12^+bsk^DN^//M6^-/-^* were stained with anti-Mmp1 or NimC1 antibody. Quantification of GFP clones’ size (B’, from left to right, n = 21, 21, 14) and relative NimC1^+^ area (B”, from left to right, n = 28, 20, 25) of indicated genotypes. **(C)** Eye discs bearing GFP labeled WT clones and *M6^-/-^* clones were analyzed for JNK activation, as indicated by the expression of TRE-DsRed. **(D)** Eye discs bearing GFP labeled clones of WT, *M6^-/-^*, and *M6^-/-^*+*bsk^DN^* were shown. Quantification of relative size of GFP^+^ region with indicated genotypes (D’, from left to right, n = 15, 23, 13). **(E)** Adult eyes of flies with indicated genotypes were shown, with the quantification of eye size (E’, from left to right, n = 12, 6, 8). **(F)** Cartoon showing the genetic basis of specifically mutating *puc* surrounding *Ras^V12^* tumors induced by the modified Q-MARCM system. **(G)** Eye-antennal discs bearing clones of *QRas^V12^//WT* and *QRas^V12^//puc^-/-^*were stained with anti-Mmp1 or NimC1 antibody. **(H)** A working model elucidating the mechanisms by which the subclonal cooperation between *M6^-/-^* and *Ras^V12^* clones induces tumorigenesis. Statistical analysis by Tukey’s multiple comparisons test (B’, D’, E’) or Dunn’s multiple comparisons test (B”); mean ± SD. ns, not significant; \*\*\*\**p* < 0.0001. Scale bars: 100μm (B, C, D, E), 50μm (G).

It is worth noting that despite the complete suppression of autonomous JNK activation, we observed a non-autonomous upregulation of Mmp1 when specifically inhibiting JNK in the *Ras^V12^* subclones of the *Ras^V12^//M6^-/-^* tumor (Fig. 4B, arrows). This finding is significant as the expression of *Ras^V12^* alone did not sufficiently upregulate Mmp1 expression (Fig. 4B). Given that JNK activity can propagate at a distance to neighboring cells (*21, 55*), we hypothesized that *M6^-/-^*cells initially trigger JNK activation, which then further propagates to adjacent *Ras^V12^* cells within the *Ras^V12^//M6^-/-^* tumor. Of note, scRNA-seq analysis of unique subclusters of non-GFP cells derived from *Ras^V12^//M6^-/-^*tumors also revealed a significant enrichment of MAPK pathway and activation of JNK signaling (Figs. 4A and S4B). Consistently, mutation or depletion of *M6* alone induced robust JNK activation in both the eye and wing epithelia, as visualized by TRE-DsRed reporter (Fig. 4C), the expression of *puckered-lacZ* (*puc-lacZ*) (Fig. S4C), and Mmp1 staining (Fig. S4D). Furthermore, in line with the essential role of JNK activation in *M6*-depleted cells, we observed that inhibition of JNK completely rescued the reduction in clone size and adult eye size, as well as the induction of apoptosis caused by the depletion of *M6* (Figs. 4D-E’, S4E, and S4E’). Finally, we genetically activated JNK in *Ras^V12^* surrounding cells by removing *puc*, the phosphatase that negatively regulates JNK activation (Fig. 4F) (*56*), and examined whether JNK activity could propagate and promote malignant transformation of *Ras^V12^* tumors. Remarkably, the deletion of *puc* in *Ras^V12^* surrounding cells led to enhanced overgrowth and JNK activation of *Ras^V12^* tumors, as well as the increased recruitment of TAM (Figs. 4G, S4F, and S4F’).

Together, these results imply that while the JNK pathway is activated in both GFP-positive and GFP-negative subclusters of *Ras^V12^//M6^-/-^*tumor, the primary source of JNK activity may be derived from *Ras^V12^*surrounding cells. This propagated JNK activity further facilitates macrophage recruitment to regulate interclonal cooperation-induced tumor malignancy.

## Discussion

Tumor heterogeneity is extensively observed and well-documented across various types of cancer (*1, 3, 4*). The existence of diverse intercellular communication between tumors and neighboring cells is crucial for tumorigenesis and the development of effective therapeutic strategies. However, the precise mechanisms underlying *in vivo* intercellular communication during tumor progression remain largely unexplored, primarily due to a scarcity of suitable *in vivo* animal models. To address this problem, we established a *Drosophila* tumor heterogeneity model by deleting the tricellular junction protein *M6* that specifically surrounds the *Ras^V12^* tumors. By combing the powerful fly genetics with the scRNA-seq, we have discovered an unrecognized role of macrophages in regulating tumor heterogeneity and malignancy by hijacking the ancient innate immune system. The loss of *M6* leads to the activation of JNK, which subsequently propagates into *Ras^V12^*tumors, thereby increasing the recruitment of macrophages. These TAMs then secrete the ligand Spz, which binds to the Toll receptor and selectively activates the Toll pathway within the *Ras^V12^* tumors. Ultimately, Toll activation synergizes with *Ras^V12^* to drive its malignant transformation by inactivating the Hippo pathway (Fig. 4H).

Our data here reveals a complex intercellular crosstalk both within epithelial cells bearing distinct oncogenic mutations and between tumors and the macrophages. Our study highlights the tumor-promoting role of Toll pathway activation in the presence of oncogenic *Ras* mutation. Apart from Toll receptor (Tl), the *Drosophila* genome encodes eight additional Toll-like receptors (TLRs). The role of different TLRs in tumorigenesis is multifaceted and can have both pro- and anti-tumorigenic effects both in fly and human, depending on the specific TLR and the context (*57*). For instance, the activation of Toll signaling hinders the elimination of polarity deficient cells and promotes their tumorous growth(*58*). Similarly, the overexpression of Toll-7 and Toll-9 induces tumor overgrowth and undead apoptosis-induced proliferation, respectively (*59, 60*). On the contrary, Toll activation in the fat body can upregulate AMP expression, leading to the remote triggering of tumor cell death (*54, 61*). Moreover, we have previously demonstrated that Toll-6 activation facilitates the elimination of precancerous *scrib* clones by inducing mechanical tension-mediated Hippo activation (*62*).

*Drosophila* is becoming an increasingly attractive model for studying questions related to human cancer biology, primarily due to the availability of sophisticated genetic tools, the high level of genetic conservation, the short life cycle, and its potential for *in vivo* testing of anti-cancer drugs (*15, 16, 41, 63–65*). Our study here emphasizes the profound significance of utilizing *Drosophila* as a model organism to study tumor heterogeneity and the intricate *in vivo* communication networks that exist between different cell types within the TME. The successful use of *Drosophila* models in uncovering conserved molecular mechanisms of tumor progression makes it intriguing to consider the potential of our fly tumor heterogeneity model to provide insights into tumor heterogeneity in humans. This, in turn, has the potential to contribute to the development of therapeutic strategies for cancer patients.

## Acknowledgements

We thank Tian Xu, Tatsushi Igaki, Duojia Pan, Lei Xue, Shian Wu, Istvan Andó, Helena Richardson, Bruce Hay, Bloomington *Drosophila* Stock Center, Vienna *Drosophila* Resource Center, and Developmental Studies Hybridoma Bank for providing fly stocks and reagents; the Microscopy Core Facility and the High-Performance Computing Center of Westlake University for the facility support and technical assistance; Wenhan Liu for fly stock maintenance. This project was supported by startup funds from Westlake University, National Natural Science Foundation of China (32170824, 32322027), HRHI program (1011103360222B1) of Westlake Laboratory of Life Sciences and Biomedicine, "Pioneer" and "Leading Goose" R&D Program of Zhejiang (2024SSYS0034).

## Author Contributions

X.M. conceived and designed the study; D.K. initiated the project and established the tumor heterogeneity model; X.M., S.Z., Y.G., and Q.X. designed the experiments and analyzed the data; S.Z. performed fly related experiments with the help from X.K., X.L., P.L., and C.W.; Y.G. performed bulk RNA-seq and scRNA-seq analyses; X.M. wrote the manuscript with input from S.Z., Y.G., and D.K.

## Competing interests

The authors declare no competing interests.

**Supplementary Materials**

Materials and Methods

Figs. S1 to S4

Tables S1 to S2

Supplemental References (66–80)

