## Supplemental Figures and legends for "Macrophages facilitate interclonal cooperation-induced tumor heterogeneity and malignancy by activating the innate immune signaling"

**Supplemental Figures and Figure legends**

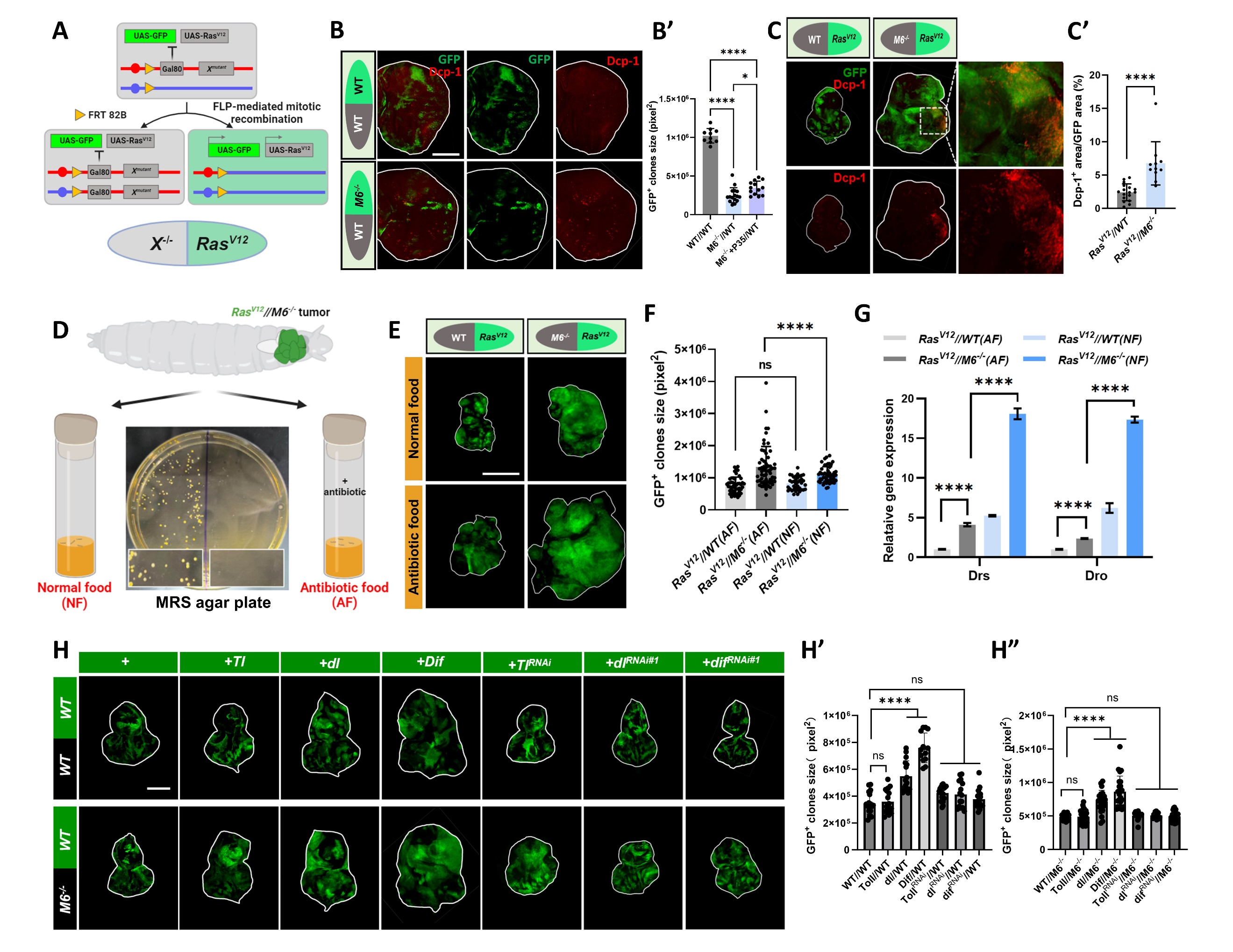

**Fig S1. *Ras^V12^//M6^-/-^* tumor-induced Toll activation is independent of microbial infection.**

(A) A cartoon is shown illustrating the genetic basis of the mutation of gene X (on chromosome 3R, as an example) surrounding the *Ras^V12^* benign tumors using the modified version of MARCM system.
(B) Eye discs bearing GFP labeled clones of *WT* and *M6^-/-^* were stained with anti-Dcp-1 antibody. Quantification of GFP^+^ clones’ size with indicated genotypes (B’, n = 10, 16, 13).
(C) Eye-antennal discs bearing clones of *Ras^V12^//WT* and *Ras^V12^*//*M6^-/-^* were stained with anti-Dcp-1 antibody. Quantification of relative Dcp-1 intensity (C’, n = 17, 11).
(D) *Ras^V12^*//*M6^-/-^* tumor bearing larvae were cultured in normal food (NF) and food supplemented with antibiotics (AF), respectively. Representative bacterial culture images were shown with larvae grown either on normal food or antibiotic food on MRS agar plates.
(E) Eye-antennal discs bearing clones of *Ras^V12^//WT* or *Ras^V12^//M6^-/-^* cultured with or without antibiotic treatment.
(F) Quantification of GFP^+^ clones’ size with the indicated genotypes (from left to right, n = 38, 47, 50, 59).
(G) qPCR analysis to determine relative *Drs* and *Dro* mRNA level of eye-antennal disc dissected from *Ras^V12^*//*WT* (AF and NF) or *Ras^V12^//M6^-/-^* (AF and NF)*.*
(H) Eye-antennal discs bearing *WT//WT*, *Tl//WT*, *dl//WT*, *Dif//WT*, *Tl^RNAi^//WT*, *dl^RNAi#1^//WT*, *Dif^RNAi#1^//WT*, *WT//M6^-/-^*, *Tl//M6^-/-^*, *dl//M6^-/-^*, *Dif//M6^-/-^*, *Tl^RNAi^//M6^-/-^*, *dl^RNAi#1^//M6^-/-^*, *Dif^RNAi#1^//M6^-/-^*. Quantification of GFP^+^ clones’ size for the indicated genotypes (H’, from left to right, n = 17, 17, 16, 13, 18, 15, 17) and (H”, from left to right, n = 18, 34, 21, 24, 13, 13, 24).

Statistical analysis by unpaired two-tailed Student's t-test (C’), Tukey's multiple comparisons test (B’, F’), or Šídák's multiple comparisons test (G); mean ± SD. ns., not significant; **p*<0.05, *****p*<0.0001. Scale bars: 100μm (B), 200μm (C, H), 400μm (E).

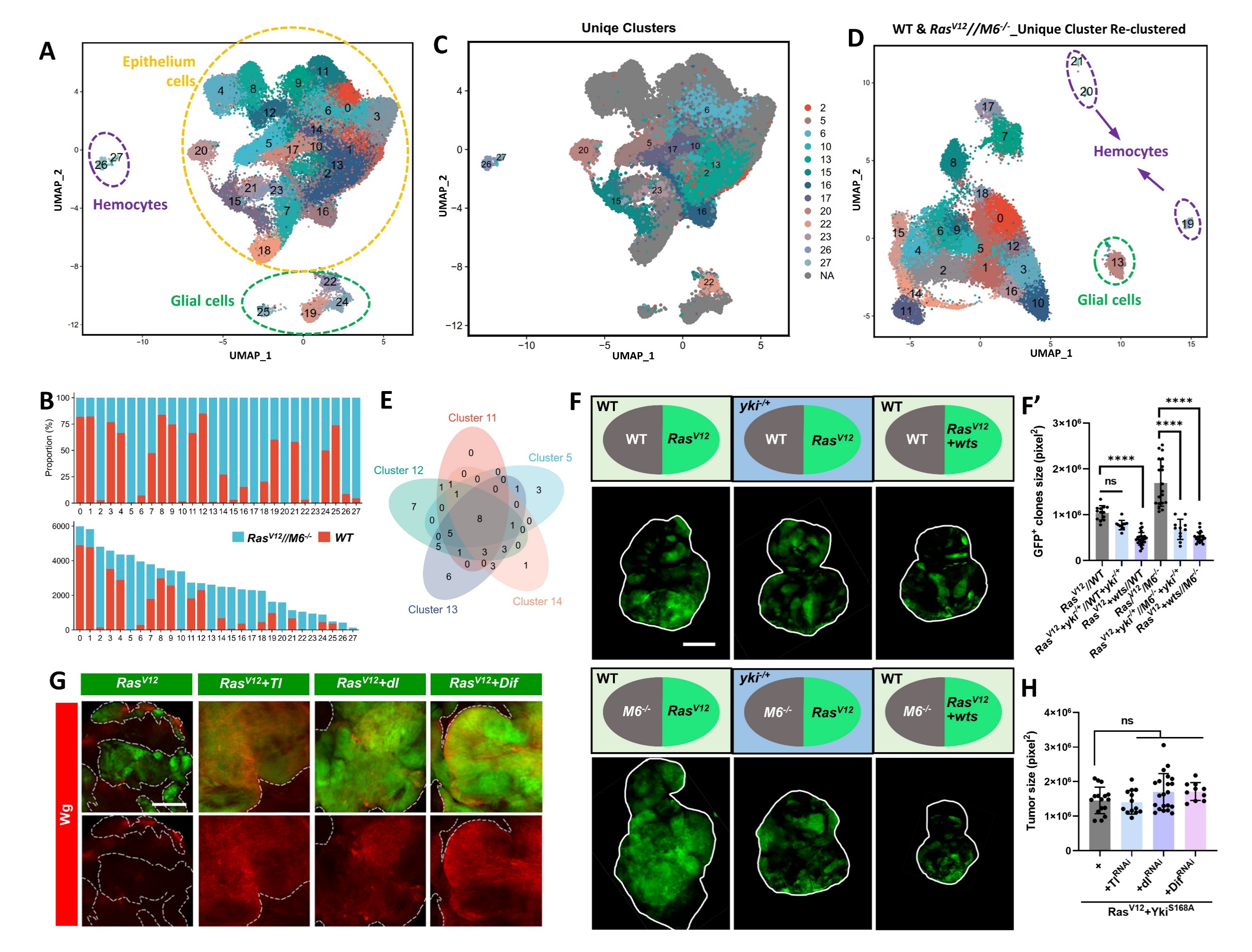

**Fig S2. Hippo signaling activation inhibits *Ras^V12^//M6^-/-^*-induced tumor overgrowth.**

(A) UMAP plot showing integrated WT and *Ras^V12^//M6^-/-^* single-cell datasets. A total of 32225 cells from WT datasets and 43992 cells from *Ras^V12^//M6^‑/-^* datasets were integrated using 2000 anchor genes. The cell type was annotated using known cell markers. Clusters 26 and 27 were Drosophila hemocytes (macrophages), clusters 19, 22, 24, and 25 were glial cells (perineurial glia, wrapping glia, and sub-perineural glia), and the remaining clusters consisted of a mixture of photoreceptors, ommatidial and interommatidial cells, undifferentiated cells, etc., which collectively form the pseudostratified columnar epithelium of the eye-antennal disc.
(B) Proportion (top) and number (bottom) of cells of each cluster from *WT* or *Ras^V12^//M6^-/-^* discs.
(C) UMAP plot showing unique clusters of *Ras^V12^//M6^-/-^* datasets. Clusters with at least 120 cells and over 80% occupancy were considered as unique ones.
(D) UMAP plot showing re-clustering of the unique *Ras^V12^//M6^-/-^* clusters in (C). A total of 22 clusters were generated (resolution = 1.2), including hemocytes and glial cells.
(E) Venn diagram showing KEGG enriched terms from different *Ras^V12^*//*M6^-/-^* unique GFP^+^ clusters.
(F) Eye-antennal discs bearing clones of *Ras^V12^//WT*, *Ras^V12^+ yki^-/+^//**yki^-/+^*, *Ras^V12^+ wts//WT*, *Ras^V12^//M6^-/-^*, *Ras^V12^+ yki^-/+^//M6^-/-^+ yki^-/+^*, *Ras^V12^+ wts// M6^-/-^*. Quantification of GFP^+^ clones’ size with indicated genotypes (F’, from left to right, n = 13, 11, 26, 19, 11, 22).
(G) Eye-antennal discs bearing clones of *Ras^V12^*, *Ras^V12^+ Tl*, *Ras^V12^+ dl*, *Ras^V12^+ Dif* were stained with anti-Wg antibody.
(H) Quantification of GFP^+^ clones’ size with indicated genotypes (from left to right, n = 16, 13, 21, 10).

Statistical analysis by Tukey's multiple comparisons test (F’, H); mean ± SD. ns., not significant; *****p*<0.0001. Scale bars: 200μm (F),100μm (G).

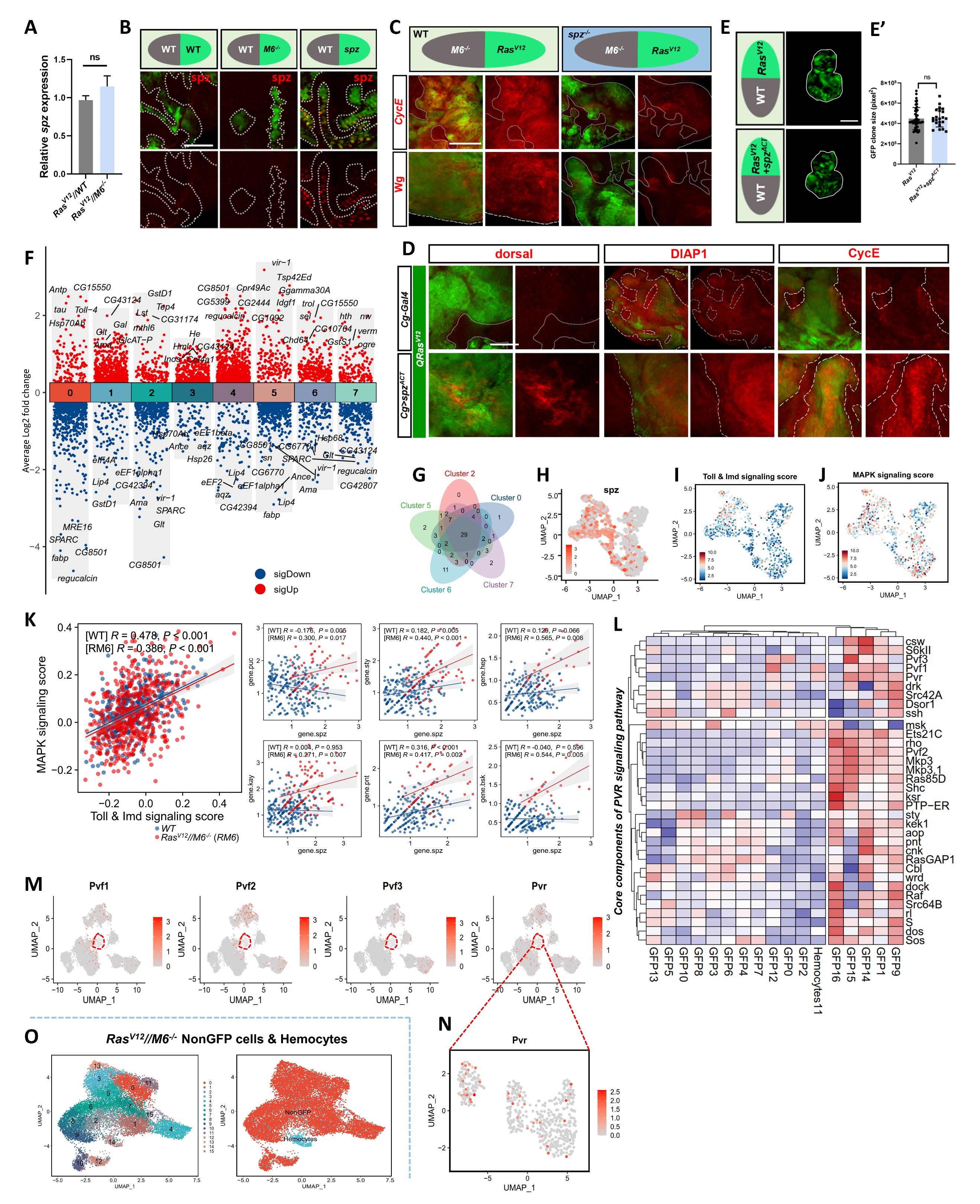

**Fig S3. The upregulation of *spz* in macrophages may be associated with Pvr-mediated JNK activation.**

(A) qPCR to determine *spz* mRNA expression of *Ras^V12^//WT* and *Ras^V12^//M6^-/-^* tumors.
(B) Eye discs bearing clones of *WT//WT*, *M6^-/-^//WT* and *spz^ACT^//WT* were stained with anti-spz antibody.
(C) Eye-antennal discs bearing clones of *Ras^V12^//M6^-/-^*, *Ras^V12^+spz^-/-^//M6^-/-^+spz^-/-^* were stained with anti-cyclin E or anti-Wg antibody.
(D) Eye discs bearing GFP labeled *QRas^V12^* clones with *spz^ACT^* overexpression specifically under the control of *Cg* promoter were stained with anti-dorsal, anti-DIAP1, or anti-Cyclin E antibody.
(E) Eye-antennal discs bearing GFP labeled clones of *Ras^V12^* and *Ras^V12^+spz^ACT^*. Quantification of GFP^+^ clone’ size with indicated genotypes (E’, from left to right, n = 50, 22)
(F) Volcano plot showing marker gene expression of each hemocyte cluster in Figure 3J.
(G) Venn diagram showing KEGG enriched terms from different *Ras^V12^//M6^-/-^* hemocytes.

(H-J) UMAP plots showing *spz* expression (H), Toll & Imd signaling score (I), and MAPK signaling score (J) of integrated WT and *Ras^V12^//M6^-/-^* hemocytes.

(K) Expression correlations analyses of WT hemocytes and *Ras^V12^//M6^-/-^* tumor-associated hemocytes. Toll & Imd signaling score and MAPK signaling are positively correlated (left), the expression of *spz* showed pan-positive correlations with MAPK signaling critical members, including *puc*, *sty*, *hep*, *kay*, *pnt*, and *bsk* (right).
(L) Heatmap of the expression patterns of the PVR signaling members of different clusters in Figure 3L. The color density reflects the average expression level of a given gene, and each row is normalized using a z-score. It is noteworthy that clusters 1, 9, 14, 15, and 16 from GFP^+^ tumors exhibited high expression of the ligands (*Pvf1*, *Pvf2*, and *Pvf3*), while hemocytes showed high expression of the receptor, *Pvr*.

(M-N) UMPA plots showing the expression of *Pvf1*, *Pvf2*, *Pvf3*, and *Pvr* across integrated *Ras^V12^*//*M6^-/-^* hemocytes and *Ras^V12^//M6^-/-^* GFP^+^ cells. (N) Zoom in UMPA plot showing the expression of *Pvr* within *Ras^V12^*//*M6^-/-^* derived hemocytes.

(O) UMPA plots showing integrated hemocytes and non-GFP cells derived from *Ras^V12^//M6^-/-^*. A total of 16 clusters were identified (left, resolution = 1.2).

Statistical analysis by unpaired t test (A, E’). mean ± SD. ns, not significant. Scale bars: 50μm (B), 100μm (C, D) and 200μm (E).

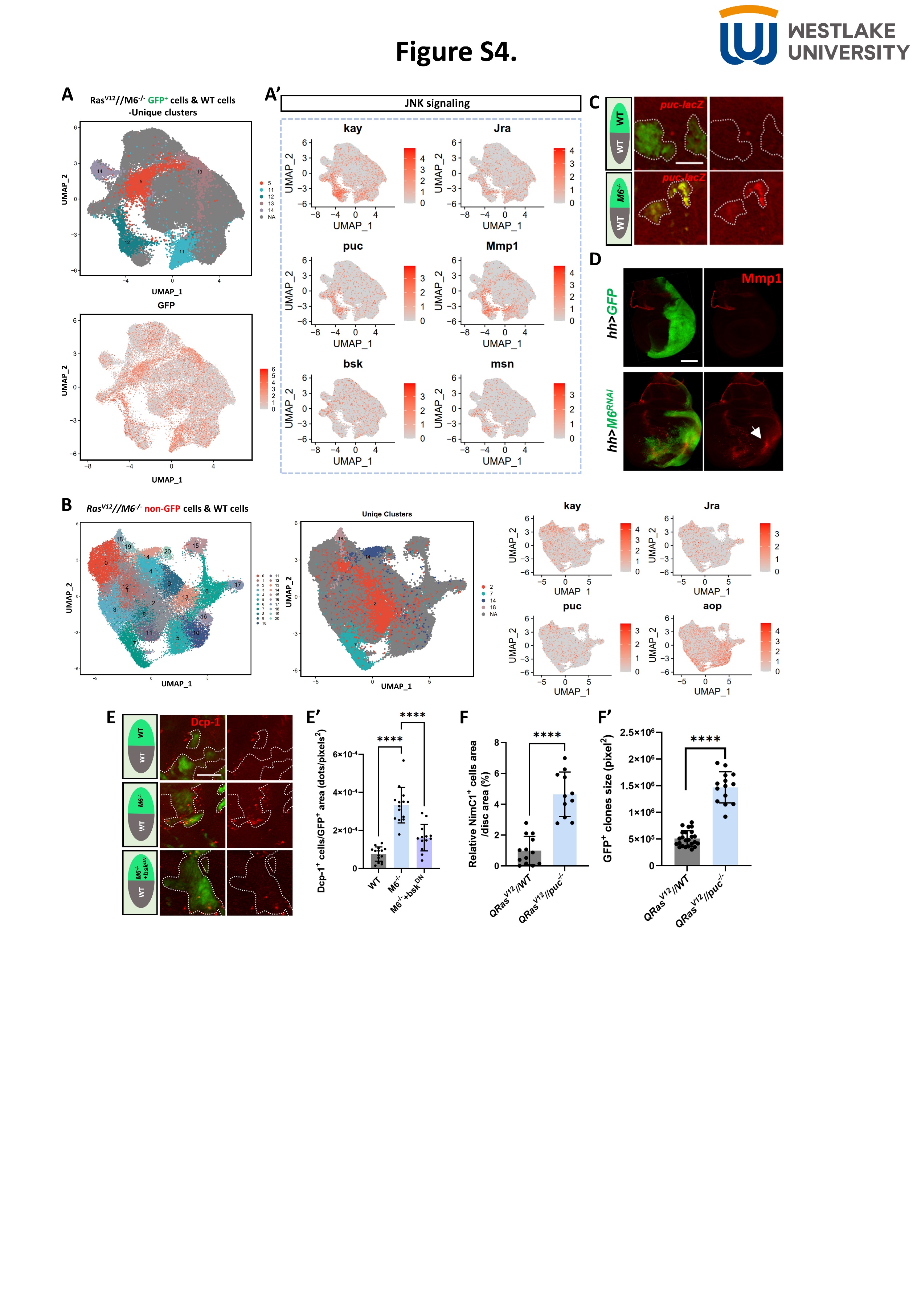

**Fig S4. Loss of *M6* activates JNK signaling and leads to cell death.**

(A) UMAP plots showing integration of *Ras^V12^//M6^-/-^* GFP^+^ cells and WT cells (resolution = 1.2). Clusters 5, 11, 12, 13, and 14 were identified as *Ras^V12^//M6^-/-^* GFP^+^ unique clusters. (A’) The expression pattern of given genes of JNK pathway enriched by KEGG analysis (*kay*, *jra*, *puc*, *Mmp1*, *bsk*, *msn*) were shown.
(B) UMAP plots showing integration of *Ras^V12^//M6^-/-^* non-GFP cells and WT cells (resolution = 1.2). Clusters 2, 7, 14, and 18 were identified as *Ras^V12^//M6^-/-^* non-GFP unique clusters (middle). The expression pattern of given genes of JNK pathway enriched by KEGG analysis (*kay*, *jra*, *puc*, *aop*) were shown (right).
(C) Eye discs bearing GFP labeled WT and *M6^-/-^* clones were stained with anti-β-galactosidase antibody to monitor *puc* transcription.
(D) Wing discs of control or with *M6^RNAi^* expression under *hh* promoter (GFP^+^ region) were stained with anti-Mmp1 antibody.
(E) Eye discs bearing GFP labeled clones of *WT*, *M6^-/-^*, and *M6^-/-^*+*bsk^DN^* were stained with anti-cleaved Dcp-1 antibody. Quantification of relative Dcp-1 intensity was shown (E’, from left to right, n = 15, 14, 13).
(F) Quantification of relative NimC1^+^ cell area (n = 13, 11) and GFP^+^ clones’ size (n=24, 14) of flies bearing *QRas^V12^//WT* and *QRas^V12^//puc^-/-^* clones.

Statistical analysis by Tukey's multiple comparisons test (E), Mann Whitney test (F) and unpaired two-tailed Student's t-test (F’); mean ± SD. *****p* < 0.0001. Scale bars: 25μm (C), 50μm (E), 100μm (D).
